# Enhancing Ligand-Based Virtual Screening with 3D Shape Similarity via a Distance-Aware Transformer Model

**DOI:** 10.1101/2023.11.17.567506

**Authors:** Manuel S. Sellner, Amr H. Mahmoud, Markus A. Lill

**Affiliations:** Department of Pharmaceutical Sciences, University of Basel, Klingelbergstrasse 50, Basel, 4056, Basel-Stadt, Switzerland; SIB Swiss Institute of Bioinformatics, Lausanne, 1015, Vaud, Switzerland

**Keywords:** Similarity search, Deep learning, Transformer model, High throughput screening, Virtual screening

## Abstract

Following the assumption that chemically similar molecules exhibit similar biologcial properties, ligand-based virtual screening can be a valuable starting point in drug discovery projects. While 2D-based similarity metrics generally focus on similar scaffolds or substructures, 3D-based methods can capture the shape of a molecule, allowing for the identification of compounds with different scaffolds. We recently published a proof-of-concept study which demonstrated how a Transformer model can be adapted to preserve 2D similarities in latent space in the form of Euclidean distances. In this work, we extend this research and prove that the approach can be adapted to 3D similarities. We use pharmacophore-based shape similarity as 3D similarity measure. We show that the model is able to enrich the predicted most similar hits with compounds with different scaffolds that are indeed similar in 3D space. Whereas classical pharmacophore- or shape-based 3D similarity methods rely on expensive alignment processes, in our approach, we identify similar compounds directly by the Euclidean distances in latent space. This enables for the first time the 3D screening of ultra-large databases with high efficiency.

## 1 Introduction

Virtual screenings of ultra-large compound libraries are receiving increasing attention in the scientific community[1–3]. While some virtual screening strategies rely on structure-based approaches [4], most use ligand-based methods due to their simplicity and computational efficiency [5–8]. In this work, we explore a novel method to accelerate 3D ligand-based ultra-large virtual screening using a Transformer-based deep neural network.

### 1.1 Similarity Search

Ligand-based similarity methods are widely used because of their computational efficiency, which is orders of magnitude faster than classical structure-based virtual screening methods. Ligand-based similarity concepts assume that chemically similar molecules exhibit similar biological activity [9, 10]. Common 2D similarity search consists of the extraction and comparison of molecular features. Usually, these features are stored in binary vectors (fingerprints) which can be easily compared using, e.g., the Tversky index, Tanimoto, or Dice coefficient. Due to the simplicity of this approach, these methods are usually fast enough to allow the screening of more than a billion compounds in a matter of hours.

On the other hand, 3D similarity methods are usually more computationally demanding. While alignment-free methods may still be computationally feasible [11], the usually more accurate alignment-based methods, such as pharmacophore or shape screening, come with an increased computational cost [12]. Nevertheless, it has been shown that even these computationally demanding methods can be used to screen billions of compounds with the right hardware. For example, Michino et al. screened approximately 1.12 billion compounds in 19.5 hours using 216 GPUs [13]. Given that not everyone has access to such significant computational resources, we believe it is essential to accelerate these accurate yet comparatively slow ligand-based screening methods.

### 1.2 Representation Learning

In the field of similarity search, a recurring issue is the maximization of the information content in a molecular fingerprint [14, 15]. Since in this work we focus on the use of deep neural networks, this problem falls into the domain of representation learning. Representation learning aims to transform raw high-dimensional data into a reduced set of features that can be used to optimally represent the data and enable their use in downstream tasks [16, 17]. Representation learning has been used in various fields such as language processing [18], time series [19], optimization of industrial processes [20], investigation of biological sensorimotor integration [21], and molecular property prediction [22].

A common approach is contrastive representation learning, in which similar samples are trained to be close together in embedding space, while dissimilar samples should be farther apart [23]. Thus, in contrastive learning, input samples are compared to each other. This allows for the use of unsupervised learning as long as input samples can be compared with a defined similarity metric. One of the earliest contrastive loss functions was developed by Chopra et al. and is used to cluster samples of the same class in a similar location in embedding space [24]. Other important loss functions used in contrastive representation learning include the triplet loss [25], lifted structured loss [26], N-pair loss [27], and noise contrastive estimation (NCE) loss [28].

Generative representation learning is another important category of representation learning [23]. In generative representation learning, a model is trained to generate new samples (or reconstruct samples from the input). The concept is that, for a model to generate realistic samples, it must learn the fundamental structure of the data.

### 1.3 Previous Work

In a proof-of-concept study, we recently demonstrated that it is possible to train a deep neural network model to create a similarity conserving latent space [29]. We demonstrated that the latent space can be shaped in such a way that allows to use the Euclidean distance between embedded molecules as a measure of their similarity. This was done using a combination of generative and contrastive representation learning. The utilized Transformer-based model reconstructed SMILES strings that were given as input. At the same time, it used a custom similarity loss for contrastive learning of continuous molecular similarities (see Equation 2).

Using this model, it was possible to reduce the search space for virtual screening by several orders of magnitude. For simplicity, we used a simple 2D similarity measure based on Morgan fingerprints. However, due to the low computational cost of calculating 2D similarities, training such a model does not give a significant benefit over directly using the underlying similarity metric. Here, we extend this work and adapt the model to computationally expensive alignment-based 3D similarity metrics, which results in significant improvements in efficiency compared to other 3D similarity methods.

### 1.4 Challenges Going From 2D to 3D

To utilize 3D similarity metrics, the architecture of the model needs to be modified to allow for 3D structural data as input instead of 1D SMILES strings. Here, we represent molecules in the form of graphs, where atoms are nodes, and bonds are edges. This approach also allows to include 3D distance information as part of the edge featurization. A detailed description of this process can be found in Section 3.3.

Arguably, the biggest challenge is the computational cost of complex 3D similarity calculations. For the model trained on 2D similarities in our previous proof-of-concept study, it was possible to either calculate all pairwise similarities in the training set before training or using online learning by calculating the similarities on the fly during training. When using alignment-based 3D similarity metrics, it is not feasible to calculate all pairwise similarities for a large dataset. Also, online learning would be simply too slow. One way to overcome this problem is to use active learning methods. Active learning is a technique to sample from unlabeled data and choose new samples to annotate and add to the training set based on a certain algorithm in order to maximize the model’s improvement [30, 31]. There are several algorithms (acquisition functions) that are often used in active learning. Regardless of the specific algorithm, their goal is always to select the best data to learn from in order to boost the model’s performance as efficiently as possible. The use of active learning therefore allows to start training on a small training set which is iteratively grown based on the selection of the implemented algorithm(s). In our case, this has the advantage that only a small portion of the data has to be annotated (i.e. similarities have to be calculated) before the training. Each active learning cycle only adds new samples that are beneficial for the model’s training, thus making the whole training process more efficient.

One commonly used active learning acquisition function is called query by committee (QBC). This algorithm employs a committee of models (so-called students). New samples for annotation are selected based on the maximum disagreement in prediction between the student models [32–34]. Therefore, a normal QBC algorithm requires multiple models to be trained in parallel. Since this comes with an additional computational cost, a committee can also be simulated by using the same model with activated dropout for the predictions. This method is also called query by dropout committee [35].

Another popular active learning algorithm is expected model change maximization (EMC). In this algorithm, the gradient of the loss with respect to an input sample is used to estimate the expected change of the model when learning from the sample [36, 37]. In practice, a set of unlabeled samples is passed through the model. For each sample, the gradient of the loss with respect to the input is calculated and the samples with the largest gradient are chosen for annotation. Since it is necessary to calculate the loss for unlabeled samples, this method cannot be used for loss functions that require labels.

### 1.5 Our Contribution

In this work, we extend our previous proof-of-concept study using 2D similarities to the use of 3D similarity metrics for efficient high-content virtual screening. We present the necessary modifications to the model architecture and the training process to enable the training on computationally expensive alignment-based similarity metrics; here shape screening implemented within the Schrödinger software suite. We also show that our model is indeed capable of conserving 3D similarities in latent space and that it can be used to efficiently identify compounds with similar 3D features. In our opinion, such a model can be very valuable in the early stages of hit identification, where the main focus is the reduction of the search space.

## 2 Results and Discussion

There are several performance criteria that our model must meet. First, since the intended use for this model is ligand-based virtual screening, it should be able to actually predict similar compounds (according to the underlying similarity metric) within the top-ranked predictions. Second, the model should actually capture 3D features and not rely solely on 2D similarities. This means that the model should be able to identify similar compounds with different chemical scaffolds. Finally, the model should prove its usefulness in a “real-world example” such as successfully reproducing known binders to a given target protein based on a reference molecule.

### 2.1 General Analysis

With this general analysis, we tested the model’s ability to find similar compounds and capture 3D information. We did this by screening several query molecules against a database of structurally diverse compounds and comparing the identified hits with the hits from the pharmacophore-based 3D shape screening. This test had two desired outcomes: 1) the model’s predictions correlate with the baseline similarities and the model is able to identify a high percentage of the top-ranked hits according to the baseline similarity method. 2) the top-ranked predictions have a high shape overlap with the query molecules.

To construct the dataset for the screening, we randomly selected a subset of approximately 50,000 compounds from the ZINC database [38]. Only compounds that were not part of the training set were selected. We then clustered the compounds using the Butina algorithm implemented in RDKit based on Tanimoto similarities based on Morgan fingerprints [39]. For clustering, we used a similarity cutoff of 0.7. In total, there were 31,856 clusters, of which only 5,234 contained more than one compound. This shows that the compounds had high structural diversity. We used the centroids of the 10 largest clusters as reference compounds for our analysis. This ensured that the screening set contained compounds with 2D structures similar to those of the reference compounds. For these reference compounds, we generated a single 3D conformer using Schrödinger’s LigPrep [40]. To create a dataset to screen, we took up to 10 compounds from the created clusters until we had a set of 10,000 compounds. For the selected 10,000 compounds, we then created up to 5 conformers each using Schrödinger’s ConfGen [41, 42]. This resulted in a total of 49,495 structures to screen. To create our baseline, we used Schrödinger’s pharmacophore-based GPU shape screening tool to screen the 10 selected references against the created dataset [12, 43]. Our model was used to screen the same 10 query molecules against the same dataset. Because there were multiple conformations per compound, the best score (highest similarity or shortest distance in latent space) was used for both methods. Table 1 shows an overview of the model’s performance. The mean Pearson correlation coefficient (PCC) was − 0.73 ± 0.13. Note that the correlation should be negative because the model predicts distances and not similarities. Thus, the smaller the predicted distance, the higher the estimated similarity. Figure 1 shows the correlation between the predicted distances and the calculated shape similarities for A) ZINC001763434742 (one of the best performing queries) and B) ZINC001183157671 (one of the worst performing queries). For ZINC001763434742, the true similarities are mostly in the range from 0.15 to 0.65 while the similarities for ZINC001183157671 are mainly in a comparable small range from 0.15 to 0.5. Thus, there seem to be no compounds that are highly similar to the query in 1B). This small range in similarity values contributes to the rather low correlation coefficient.

**Table 1:** Performance analysis for 10 query molecules screened against approximately 10,000 compounds from the ZINC database. “Mean similarity top 100” shows the mean shape similarity of the top 100 calculated hits according to the baseline method. “Mean similarity top 100 pred” shows the mean predicted similarity of the top 100 predicted hits according to the model. Since the model predicts distances and not similarities, the predicted similarity *ŝ* was calculated as 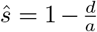 where *a* is the scaling factor as in Equation 2 and *d* is the predicted latent space distance.

| Query | PCC | Precision top 100 | Precision top 1000 | Mean similarity top 100 | Mean similarity top 100 pred |
| --- | --- | --- | --- | --- | --- |
| ZINC000570771518 | -0.78 | 0.17 | 0.47 | 0.54 | 0.46 |
| ZINC000950159323 | -0.77 | 0.30 | 0.47 | 0.56 | 0.49 |
| ZINC000954430177 | -0.54 | 0.16 | 0.33 | 0.54 | 0.44 |
| ZINC000970035445 | -0.79 | 0.33 | 0.53 | 0.57 | 0.51 |
| ZINC001183157671 | -0.64 | 0.14 | 0.35 | 0.44 | 0.36 |
| ZINC001281147597 | -0.75 | 0.25 | 0.49 | 0.54 | 0.48 |
| ZINC001368797027 | -0.78 | 0.20 | 0.44 | 0.49 | 0.43 |
| ZINC001711902206 | -0.84 | 0.34 | 0.69 | 0.49 | 0.45 |
| ZINC001740566933 | -0.88 | 0.30 | 0.76 | 0.54 | 0.50 |
| ZINC001763434742 | -0.87 | 0.58 | 0.75 | 0.60 | 0.57 |

**Fig. 1:**
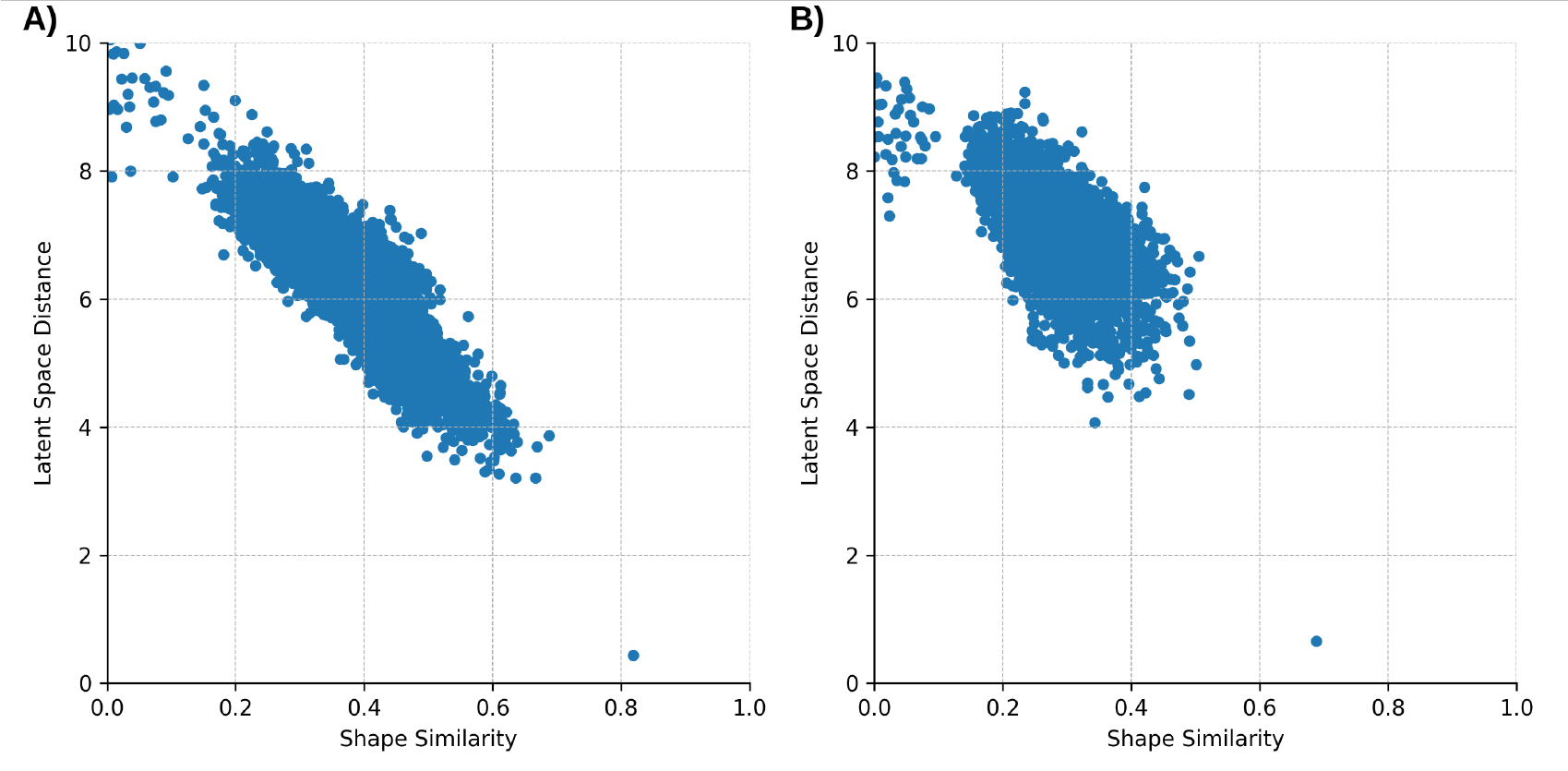
Correlation between predicted distance and calculated shape similarity for two query compounds. **A)** ZINC001763434742, *R*^2^ = 0.76. **B)** ZINC001183157671, *R*^2^ = 0.40

Using the latent space distance *d* and the scaling factor *a* used to train the model (see Equation 2), it is possible to approximate the similarity *ŝ* to 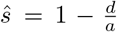. We used this equation to calculate the similarity for the top 100 predictions and compare their mean with the mean similarity of the top 100 hits from the baseline. Ideally, these two means are the same, as this would indicate that the model was able to reproduce the true similarity values. According to Table 1, the difference between these values ranged from 0.03 to 0.10, indicating that the performance depends on the chosen query compound. It can also be seen that it correlates nicely with the PCC. The precision shows the fraction of the top N predicted hits that are actually among the top N according to the baseline method. For *N* = 100, these values were generally quite low, indicating that the model was not very good at reproducing the top 100 hits among the top 100 predictions. However, the values seem to correlate with the overall performance for the specific queries, as indicated by the PCC. With some exceptions, these values are better for queries with a higher similarity to the top hits (represented by the mean similarity of the top 100 hits). This indicates that the model may generally perform better when there are compounds in a database that are highly similar to the query. Since reproducing the top 100 hits is a very difficult task and we did not expect the model to actually excel at it, we also calculated the precision for the top 1000 hits. There, the performance is generally higher, but varies greatly between the different queries, and again seems to correlate quite nicely with the PCC. On average, 53% ± 15% of the top 1000 predicted compounds were actually among the top 1000 hits according to shape screening.

To assess the screening performance of the model in more detail, we calculated the receiver operating characteristics (ROC) curves and enrichment factors (EF) for the 10 queries in Table 1. To calculate the ROC curves, we defined the first 100 hits from the shape screening as active and the rest as inactive. The goal of the model should be to accurately replicate the 100 hits as early as possible. The results of this analysis are shown in Table 2 and detailed ROC curves and reproduction plots can be found in the Supporting Information in Figures S1-10. The average area under the ROC curve was 0.93 ± 0.04, indicating a very good screening performance. Also, the mean 1% EF was 27.5 ± 13.0. This means that on average the model was able to reproduce 27.5 times more active compounds than random selection when considering only the 1% top ranked predictions.

**Table 2:** Screening performance for 10 query molecules screened against approximately 10,000 compounds from the ZINC database. The area under the receiver operating characteristics curve (AUROC) and the enrichment factors (EF) at 1, 2, 5, and 10 percent are shown.

| Query | AUROC | EF 1% | EF 2% | EF 5% | EF 10% |
| --- | --- | --- | --- | --- | --- |
| ZINC000570771518 | 0.92 | 17.0 | 14.0 | 10.0 | 6.4 |
| ZINC000950159323 | 0.95 | 29.0 | 21.5 | 13.4 | 8.0 |
| ZINC000954430177 | 0.88 | 15.0 | 11.0 | 9.0 | 6.1 |
| ZINC000970035445 | 0.96 | 33.0 | 26.0 | 15.0 | 8.8 |
| ZINC001183157671 | 0.86 | 14.0 | 13.0 | 8.0 | 5.2 |
| ZINC001281147597 | 0.94 | 25.0 | 18.5 | 13.0 | 7.9 |
| ZINC001368797027 | 0.92 | 20.0 | 15.0 | 10.6 | 7.0 |
| ZINC001711902206 | 0.96 | 34.0 | 25.5 | 14.4 | 9.4 |
| ZINC001740566933 | 0.98 | 30.0 | 27.0 | 17.0 | 9.9 |
| ZINC001763434742 | 0.99 | 58.0 | 38.5 | 19.6 | 10.0 |

To get a full picture of the performance of the model, it is important to analyze examples in which the model predicted very wrong values. Therefore, we picked examples that had a short predicted distance while having a low calculated shape similarity. Figure 2 shows one of the cases where there is a large discrepancy between the predicted and calculated rank of the compound. Although the model ranked this compound 3680 ranks too high, the shape overlap with the query molecule seems to be high. In this case, the calculated similarity may be reduced due to few matching pharmacophores. This would indicate that the model has problems learning the pharmacophore information while being able to capture the 3D shape information well. Indeed, while the pharmacophore-based shape similarity between the two molecules was 0.35, the shape-only similarity (without pharmacophore information) was 0.63.

**Fig. 2:**
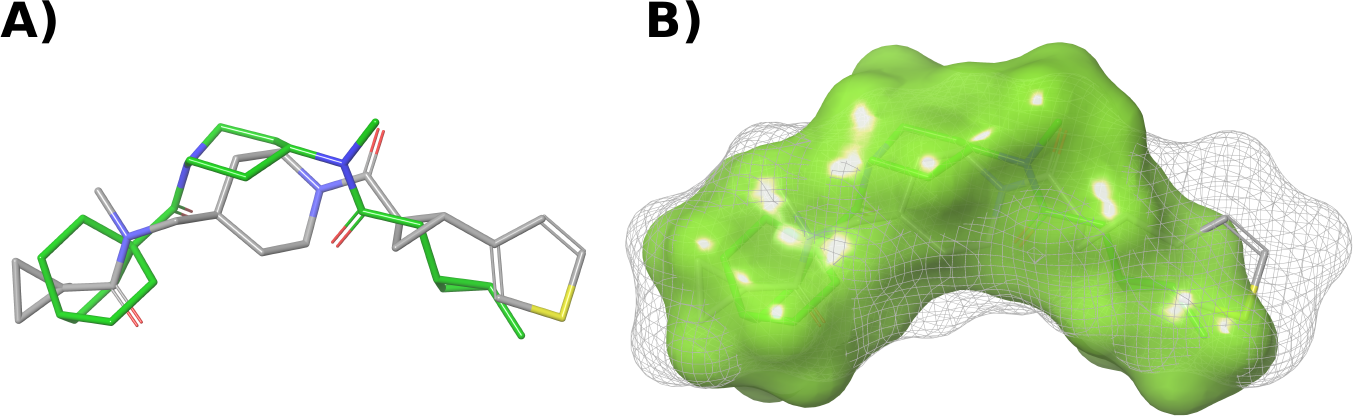
Shape overlap between ZINC000954430177 (query, green, solid surface) and ZINC001497961236 (grey, mesh surface) which was ranked top 8 by the model and rank 3688 by the baseline.

Another example in which the model overestimated a compound by 1571 ranks is given in Figure 3. It can again be seen that the overlap between the two compounds is rather high. In this example, the 2D similarity between the two compounds is low (0.35) and it can be seen that the predicted similar compound has a scaffold different from the query. Instinctively, one may say that this is actually a good hit, but nevertheless the model did not reproduce the correct rank as calculated by the baseline. Comparing the pharmacophore-based shape similarity (0.41) with the shape-only similarity (0.63) shows again that the discrepancy of the ranks was caused by the model’s inability to capture the pharmacophore information while nicely reproducing the 3D shape overlap.

**Fig. 3:**
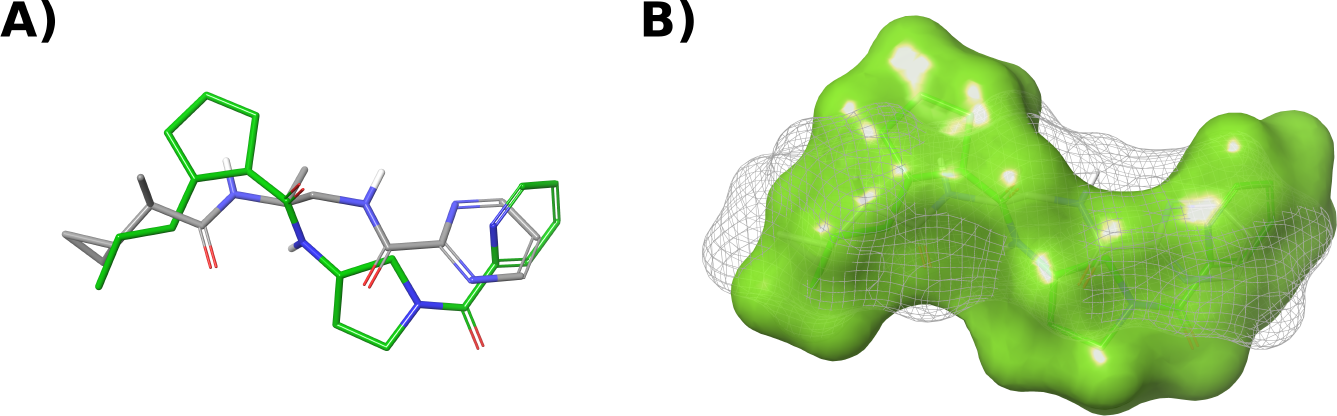
Shape overlap between ZINC000570771518 (query, green, solid surface) and ZINC001386435162 (grey, mesh surface) which was ranked top 14 by the model and rank 1585 by the baseline.

There are, however, also instances where the model very nicely reproduced hits, even if the 2D structure was very different from the query. One such example is shown in Figure 4. In this case, the shape overlap appears to be worse than in the previous examples, even though this compound was highly rated by the baseline. The shape similarities with and without pharmacophore information were very similar at 0.58 and 0.59, respectively. This indicates that the score was mainly influenced by the 3D shape and that the pharmacophore information did not contribute much. Since our model performed very well on this example, it again suggests that the model is good at finding compounds with a similar shape, but not as good at capturing pharmacophore information.

**Fig. 4:**
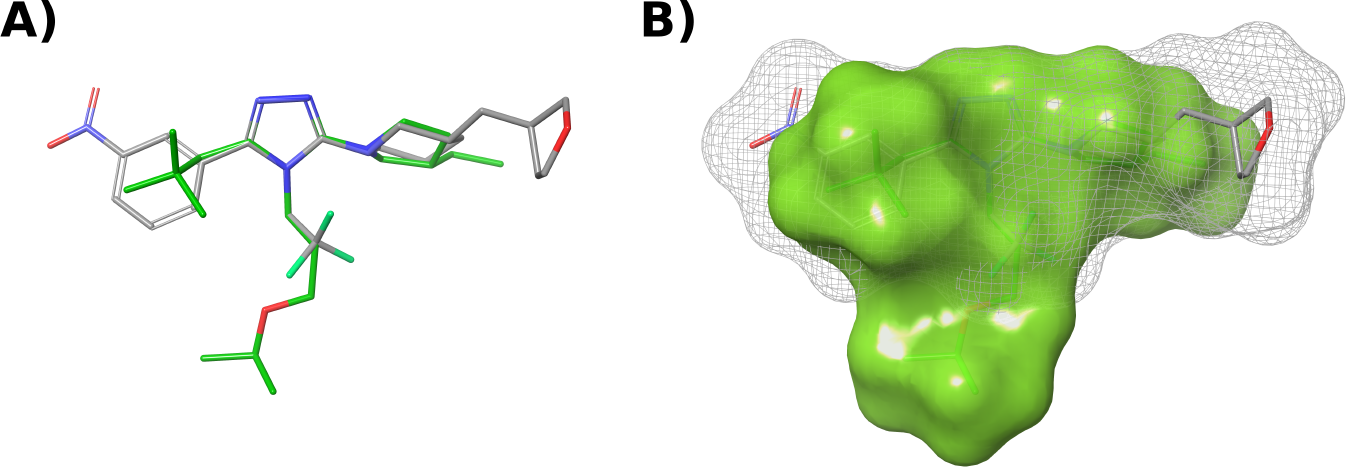
Shape overlap between ZINC001763434742 (query, green, solid surface) and ZINC001556038399 (grey, mesh surface) which was ranked top 34 by the model and rank 76 by the baseline.

In these experiments, we could show that the model is able to produce hits with a high overlap with the query molecules in 3D space. However, there may still be some shortcomings in certain cases in reproducing the exact similarity metric, especially if the similarity goes beyond “simple” 3D overlap. Nevertheless, the model proved to be able to capture 3D similarities independent of the 2D structure of the molecules.

### 2.2 Real-World Examples

To simulate a real-world example of a possible screening study, we selected 2 co-crystallized ligands as queries to screen the drugs contained in the Drugbank [44]. In a first trial, we selected raloxifene (co-crystallized to the estrogen receptor (ER) in PDB ID 1ERR). This compound has not been seen by the model before. The precision of the top 100 predictions was 53%, which is much higher than for most of the examples in Subsection 2.1. However, the precision of the top 1000 predictions was slightly lower at 49%. The PCC of all predicted distances with calculated similarities was -0.77. Interestingly, the model was able to reproduce the top 10 most similar compounds according to the baseline within the top 47 ranked hits. Since we wanted to know if the model can be used to find other compounds that modulate the ER, we investigated the top 10 predictions. Table 3 shows the results of the analysis. All 10 predicted most similar compounds have literature confirmation of ER modulating activity. While this is a very promising result, one also needs to keep in mind that the Drugbank is a biased database in that it contains not only mostly drug-like molecules, but also many known ER modulators.

**Table 3:**
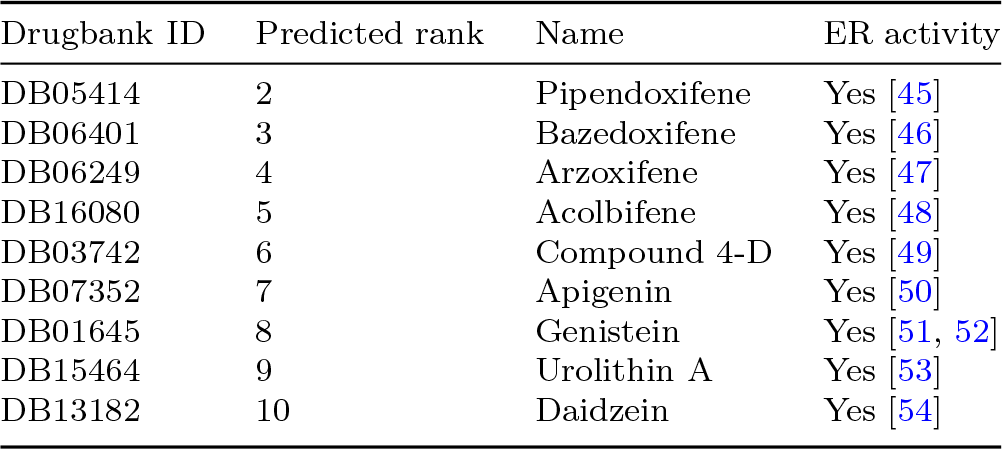
Predited top 10 most similar compounds to raloxifene. The first hit is omitted from the table because it is raloxifene itself.

| Drugbank ID | Predicted rank | Name | ER activity |
| --- | --- | --- | --- |
| DB05414 | 2 | Pipendoxifene | Yes [45] |
| DB06401 | 3 | Bazedoxifene | Yes [46] |
| DB06249 | 4 | Arzoxifene | Yes [47] |
| DB16080 | 5 | Acolbifene | Yes [48] |
| DB03742 | 6 | Compound 4-D | Yes [49] |
| DB07352 | 7 | Apigenin | Yes [50] |
| DB01645 | 8 | Genistein | Yes [51, 52] |
| DB15464 | 9 | Urolithin A | Yes [53] |
| DB13182 | 10 | Daidzein | Yes [54] |

Figure 5 shows the 2D structures of raloxifene and the 9 most similar compounds predicted by the model. It also contains the 3D structures of selected top-ranked compounds aligned with raloxifene. We further calculated the 2D similarity between the reference and the hits. While the 5 top ranked compounds were structurally quite similar (2D similarity between 0.44 and 0.66), the next 4 hits had scaffolds very different from raloxifene. They consisted of 3 isoflavonoids and 1 benzo-coumarin. The 3D alignment shows that even the compounds with different scaffolds share similar features, which could reproduce the binding mode of raloxifene. This analysis clearly shows that the model is able to find structurally dissimilar compounds by learning 3D features.

**Fig. 5:**
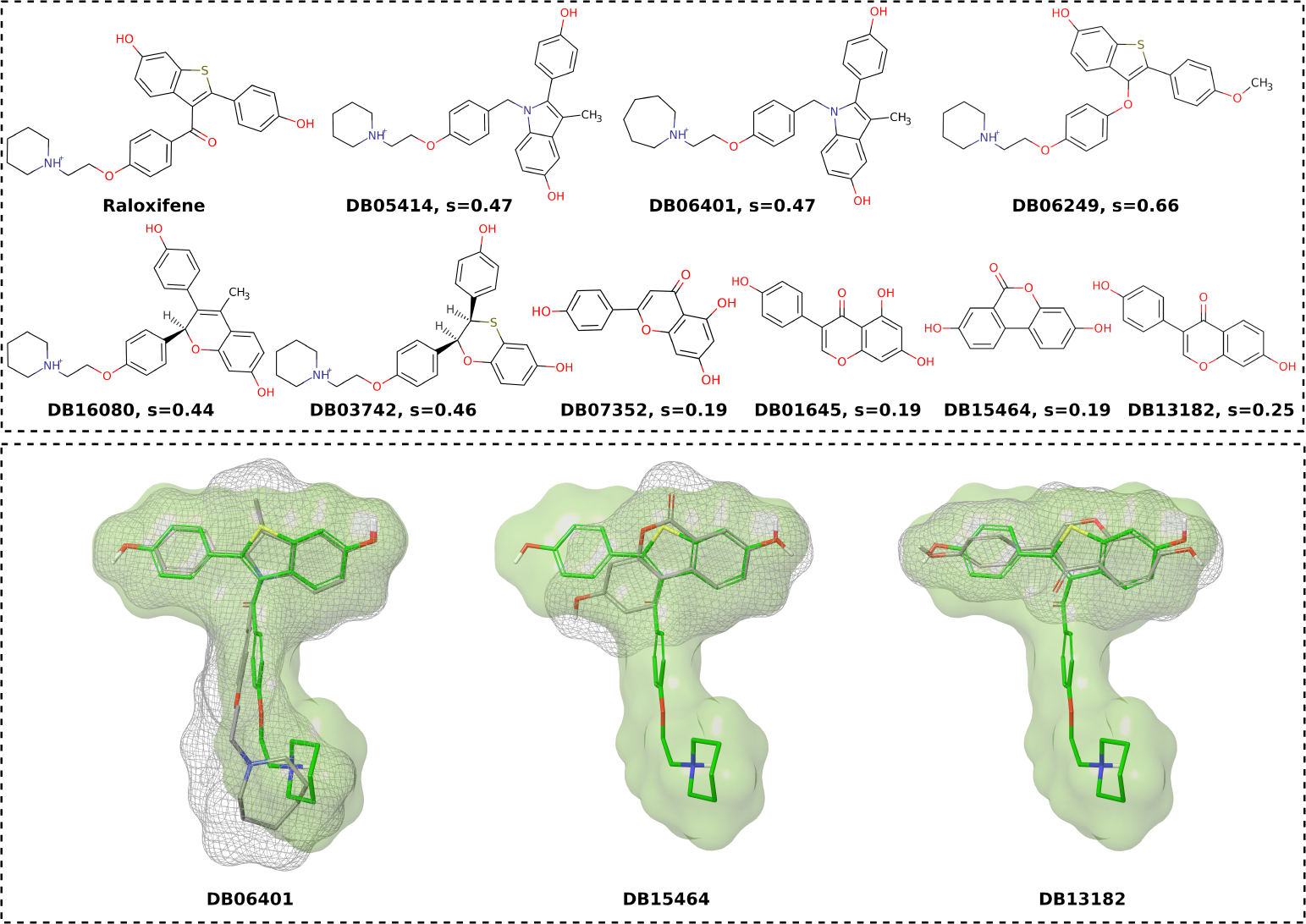
2D and 3D structures of the 9 predicted most similar compounds to raloxifene. Top: 2D structures with 2D similarity *s*. Bottom: 3D alignment of raloxifene (green stick representation with green solid surface) with 3 of the top ranked compounds with diverse scaffolds (gray stick representation with mesh surface).

Next, we wanted to test a compound with a different chemical scaffold than raloxifene. We decided to screen tetrahydrogestrinone against the drugs contained in the Drugbank. Like before, this compound has not been seen by the model before. The 10 highest ranked compounds predicted by our model are shown in Table 4. The precision of the top 100 predictions was 72%, which is exceptionally high. We believe that this is because steroidal compounds have a very unique shape and therefore it may be easier for the model to find similar compounds. The precision of the first 1000 hits was equally high with 69% and the correlation between all predicted distances and the calculated shape similarities was −0.82. The model was able to reproduce the top 10 hits from shape screening within the top 27 predictions. Eight of the 9 hits listed in Table 4 have been shown in the literature to have androgenic or anti-androgenic activity. Nevertheless, we still acknowledge the fact that the Drugbank may be a biased database in that molecules with androgenic or anti-androgenic properties are overrepresented compared to other databases.

**Table 4:** Predited top 10 most similar compounds to tetrahydrogestrinone. The first hit is omitted from the table because it is tetrahydrogestrinone itself.

| Drugbank ID | Predicted rank | Name | AR activity |
| --- | --- | --- | --- |
| DB11619 | 2 | Gestrinone | Yes [55] |
| DB11372 | 3 | Altrenogest | Yes [56] |
| DB02998 | 4 | Metribolone | Yes [57] |
| DB13563 | 5 | Norgestrienone | Yes [58] |
| DB11551 | 6 | Trenbolone | Yes [59] |
| DB06730 | 7 | Gestodene | Yes [60] |
| DB09389 | 8 | Norgestrel | Yes [61] |
| DB13602 | 9 | Promegestone | No [62] |
| DB00367 | 10 | Levonorgestrel | Yes [63] |

Since steroidal compounds have such a distinct chemical structure (and shape), we wanted to see if the model can also find non-steroidal compounds as easily as shape screening. This would show that the model indeed learns from the provided 3D information instead of relying on 2D similarity. The first non-steroidal compound we identified in the baseline hits was borneol at rank 123. Despite the fact that borneol has a very different 2D structure than steroids, the model predicted this compound to be at rank 95. This underlines the model’s ability to capture 3D similarities in latent space.

Based on these examples, we believe that our model is indeed suitable for use in real-world virtual screening applications. Although it is not able to perfectly reproduce the similarities found in the chosen baseline, its predictions are reasonable and useful in finding compounds with similar 3D features.

### 2.3 Investigation of Computational Cost

To assess whether our method actually allows screening (ultra) large databases at reduced computational cost, we screened 27 query compounds against databases of different sizes. We chose to screen 27 query compounds to be able to directly compare the results with those of Michino et al. [13] (as introduced in Section 1.1). We employed faiss, a tool created by Meta, to scan the databases [64]. Faiss allows to create searchable indexes from vectors. Several indexes with varying levels of accuracy and speed are available. We chose IndexFlatL2, an index that computes the exact squared L2 norm between the queries and all elements in the database. This index is the most accurate, but also the slowest. Thus, it would be possible to further increase the performance of the screening by choosing other more approximating indexes.

We tested databases containing 1k, 10k, 100k, and 1M compounds for screening. A screening consists of 3 steps: encoding into latent space, generation of the index, and screening of the index. Encoding into latent space is the most time-consuming task followed by the index generation (cf. Figure 6). However, these two tasks only need to be completed once, and the encoded data and index created can be reused for any future screenings. Figure 6 shows that the screening times required increase linearly with the size of the database. This allows for easy extrapolation to larger databases. Therefore, screening 27 queries against 1 billion conformers would require roughly 7.8 hours on a single CPU core. Michino et al. screened approximately 1.12 billion compounds with 10 conformers each (i.e. 11.2 billion conformers) in 19.5 hours using 216 GPUs. Extrapolating our tests to 11.2 billion conformers would result in approximately 87 hours or 3.6 days on a single CPU core. When parallelized to 8 cores (which is very reasonable for standard consumer-grade computers), the screening could be completed within 10.9 hours. If more powerful hardware is available, for example a CPU with 64 cores, the screening could be performed in only 82 minutes. We deliberately ran all experiments on a regular desktop computer with one RTX 2080 Super GPU. The GPU was used only to encode the conformers into latent space. Thus, we show that this technology enables the screening of ultra-large databases without the need for expensive hardware.

**Fig. 6:**
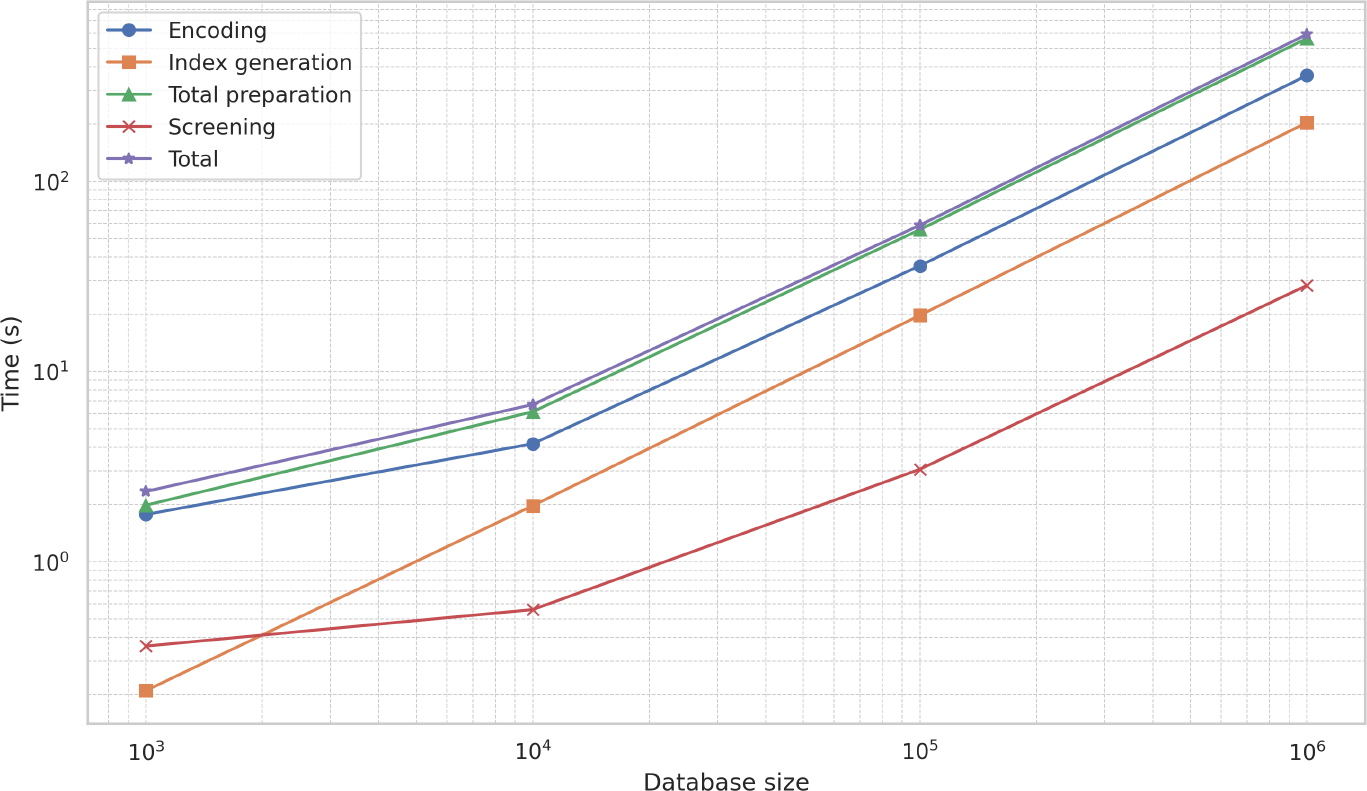
Times in seconds to screen 27 query molecules against databases of different sizes. Displayed are the time to encode the database into latent space (blue), the time to generate the IndexFlatL2 index in faiss (orange), the total preparation time (encoding plus index generation; green), the time to actually screen the database (red), and the total time (purple).

Another advantage of this method is that it allows parallel search of multiple queries. It took only an additional 2.35 seconds to screen 100 queries instead of 27 against a database containing 1 million conformers. Thus, the gain in speed over the classical shape screening increases with the number of query compounds.

## 3 Methods

### 3.1 Dataset Preparation

Like in our proof-of-concept study, we used a randomly selected subset from the ZINC database containing around 500k compounds to train our model [29]. From this dataset, we selected the 25k molecules that best cover the chemical space of the complete dataset using the Kennard-Stone algorithm [65]. From this narrowed-down subset, we again applied the Kennard-Stone algorithm to isolate the most diverse 5k compounds for use as our validation set. The remaining 20k compounds were used as the initial training set. The initially unused 475k compounds were split into an external test set, consisting of 10k compounds, and a pool of molecules that were used as an unlabeled set to sample from during the active learning cycles.

### 3.2 Data Handling

In this work, the model was trained to encode 3D molecules into latent space and decode them to SELFIES [66, 67]. In order to enable the model to learn from 3D structures, the first step was to calculate atom features to be used as nodes. This was done using RDKit. For each atom in a molecule, we encoded the atom type, the atom degree (i.e. the number of neighbors), the number of connected hydrogen atoms, the implicit valence, the hybridization, and the aromaticity in a vector that could later be used in a learnable embedding.

The second step in encoding 3D information was to create edges between nodes in a way that conserved the 3D topology. This was done by passing the Euclidean distance matrix of a molecule through an exponential decay function and combining the result with the adjacency matrix. This is described in Equation 1.

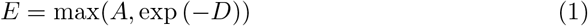

Where *A* is the adjacency- and *D* the Euclidean distance matrix of a molecule. This approach allows 3D information to be encoded in a translation and rotation invariant way while also preserving information about atom connectivity.

Given the vast amount of data and the expense of training, we chose to train the model in this initial 3D-enabled version with only one conformation per molecule. However, we think that including multiple conformations could enhance the model’s capacity to learn 3D similarities.

The SELFIES used in this work were converted from canonical SMILES which were created using Openbabel version 3.0.0 [68]. Each SELFIES that was passed through the model was tokenized based on its individual components. We decided to use SELFIES instead of SMILES strings due to their robustness.

As our baseline similarity method, we used Schrödinger’s pharmacophore-based shape similarity shipped with their 2023-2 release [43]. Unless otherwise stated, all molecules were processed with LigPrep prior to shape screening. We chose to generate protonation states at pH 7.4 and for each molecule, we created one 3D conformation using the OPLS4 force field.

### 3.3 Model Architecture

The architecture of the model had to be only minimally adapted from our proof-of-concept study. Since the 3D- and connectivity information of the molecules are fully encoded by their edges, no positional encoding is needed in the encoder. In fact, removing the positional encoding is required for permutation equivariance because the order of the nodes does not matter, and thus a positional encoding would give incorrect information. The rest of the model is still based on the original implementation of the Transformer model by Vaswani et al. [69]. In the previous study, a masked mean was used to combine the nodes to generate a latent vector, whereas this work utilizes a weighted sum pooling technique. In this method, the weights are calculated using two linear layers with a tanh activation between them.

To train the model, we used the same combination of the reconstruction (i.e. cross-entropy) loss and our custom similarity loss as in the proof-of-concept study. Equation 2 shows this loss function in detail where *A* is an anchor sample, *X* is some other sample in the mini batch, *sim*(·, ·) is a similarity function (in this case Schrödinger’s pharmacophore-based shape similarity), and *f* (·) is the encoder of the model, encoding a molecule into latent space. To decrease the density of the latent space, the scaling factor *a* is used. This scaling factor was set to 10 in this work.

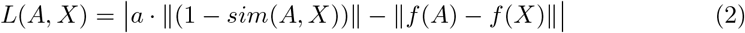

Like in our proof-of-concept study, we used scaled dot-product attention as described in Equation 3, where *Q, K*, and *V* are tensors containing the queries, keys, and values, and *d*_*k*_ is the dimensionality of the keys [69].

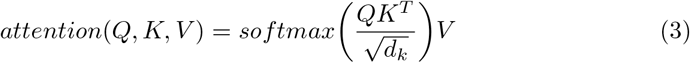

The complete model architecture is depicted in Figure 7. The model was trained with a batch size of 64, a latent space dimensionality of 252, a learning rate of 1*e*^*−*4^, and 4 encoder and decoder layers each. Each attention module consisted of 4 heads and the model was trained for a minimum of 200 and a maximum of 800 epochs per active learning cycle.

**Fig. 7:**
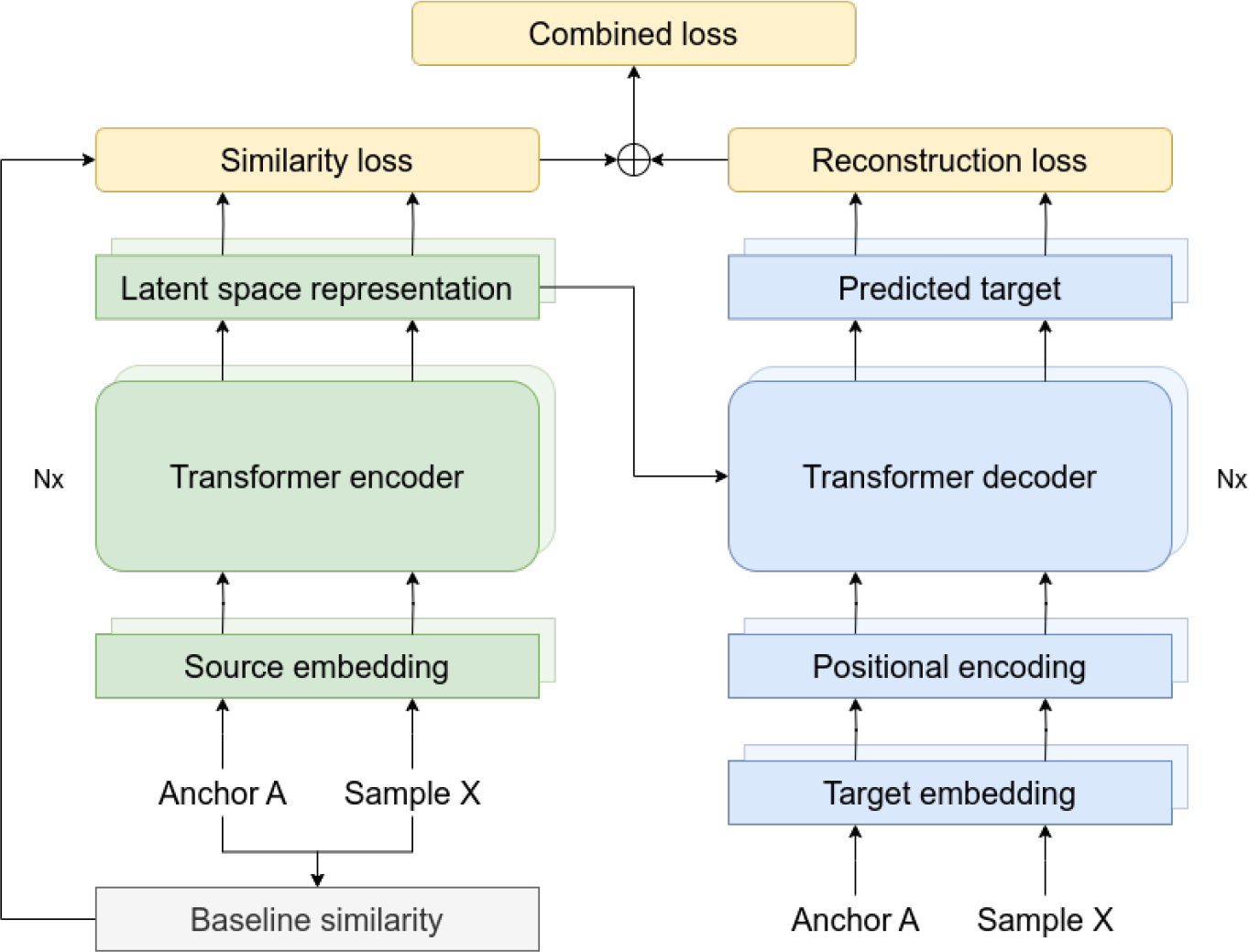
Architecture of the model used in this study. This adaptation of the original Transformer implementation enables the encoding of molecular three-dimensional information that is invariant to translations and rotations. We used Schrödinger’s pharmacophore-based shape similarity as baseline.

Because we used an active learning approach, the initial training set was much smaller compared to the training set used in the proof-of-concept study. However, from the two implemented loss terms, only the similarity loss depends on labeled data, whereas the reconstruction term is completely unsupervised. Therefore, for each mini batch, we created an additional mini batch containing random samples from the unlabeled dataset. This mini batch was used to train only the reconstruction. We chose this approach to improve the reconstruction abilities of the model while preventing overfitting on the small datasets.

Since calculating the 3D similarities is comparatively expensive, we precomputed pairwise similarities for our initial training and validation set. During training, we used a similarity sampler to ensure that each molecule in a mini batch contained at least one similar compound. This was accomplished by randomly sampling 3 compounds from the 100 most similar to a given anchor molecule and adding them to the mini batch. This step is imperative for the model to conserve similarities in latent space and has already been described in our proof-of-concept study.

### 3.4 Active Learning

In this work, we used an active learning approach using a combination of QBC and EMC acquisition functions. Calculating the EMC involves calculating the gradient of the loss with respect to the input. Thus, one needs to be able to calculate the loss of a sample. Since the point of active learning is to select samples from unlabeled data to be labeled, this approach cannot be applied to the similarity loss. However, it is possible to use EMC for the reconstruction loss. For each active learning cycle, we encoded and decoded all unlabeled samples and calculated the gradient of the reconstruction loss with respect to the input. The gradients were then normalized using the L2 norm and squared to ensure positivity. This is described in Equation 4 where ∇_*x*_*L* is the gradient of the reconstruction loss *L* with respect to the input sample *x*.

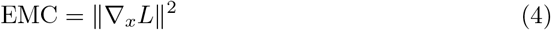

The EMC was mixed with QBC in order to also sample based on the similarity loss. To achieve this, we predicted the latent vector for each unlabeled sample 100 times using a dropout rate of 10%. We then calculated the mean over the variance of the predictions. This is shown in Equation 5 where *P* is a matrix containing *N* predictions and Var(·) is the variance.

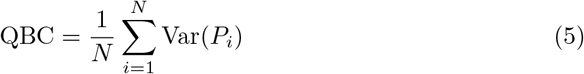

The sampling of the unlabeled data was performed on the basis of the magnitude of the EMC and QBC values. Starting from the samples with the highest values, an equal number of samples were drawn based on the EMC and QBC values. This was done until a total of 5k unique molecules were sampled from the unlabeled dataset. These samples were then labeled by calculating pairwise similarities (including the existing labeled samples) and added to the training set. Thus, for each active learning cycle, the training set grew by 5k compounds. Because we calculated pairwise similarities, the time used to label the newly sampled compounds increased exponentially. The model in this work was trained for 5 active learning cycles, resulting in a final training set containing 45k compounds.

## 4 Conclusion

We previously demonstrated that a distance-aware transformer model can be used to preserve 2D similarities in latent space. We claimed that this method can be used independent of the underlying similarity metric, allowing to efficiently estimate highly complex 3D similarities.

In this work, we show how a slightly adapted model is capable of capturing such 3D similarities. We demonstrated that our model, which uses a translation and rotation invariant molecular representation, is able to recognize 3D features of molecules and identify molecules with similar shapes in a virtual screening context. Although the model cannot perfectly reproduce the underlying pharmacophore-based shape similarity, it is still capable of enriching the top hits with highly similar compounds. In fact, we believe that using a shape-only similarity metric would lead to much better performance because the model does not seem to be able to fully capture the pharmacophore information. Thus, for such special similarity metrics, the model might need to be further adapted to better reproduce the baseline similarity.

The approach described herein enables the use of the Euclidean distance in latent space as an approximation of computationally expensive 3D similarity metrics. It therefore allows researchers to run quick and efficient (pre)screenings on ultra-large databases using regular low-cost computer hardware.

## Supplementary information

This article has an accompanying supplementary file.

## Acknowledgments

We gratefully acknowledge the support of NVIDIA Corporation with the donation of two RTX A5000 GPUs used for this research.

