## Supplementary Information for "Enhancing Ligand-Based Virtual Screening with 3D Shape Similarity via a Distance-Aware Transformer Model"

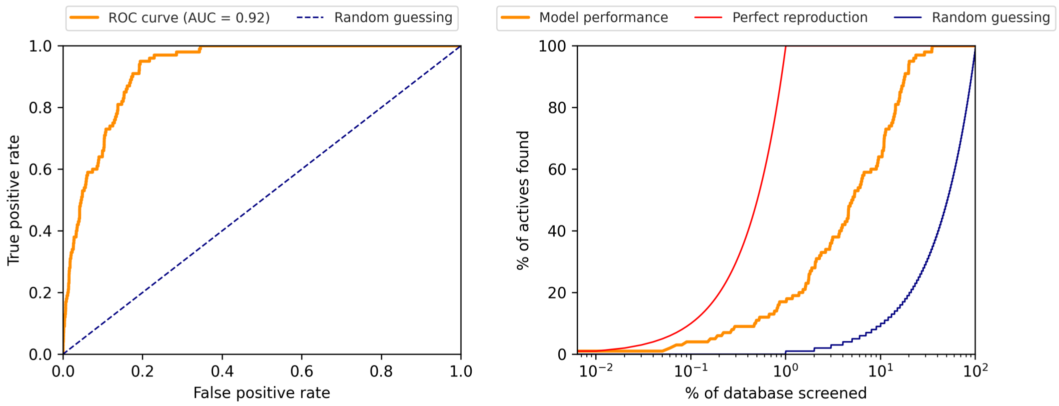


**Fig. S1**: Screening performance for query ZINC000570771518. Left: ROC curve, right: reproduction performance.

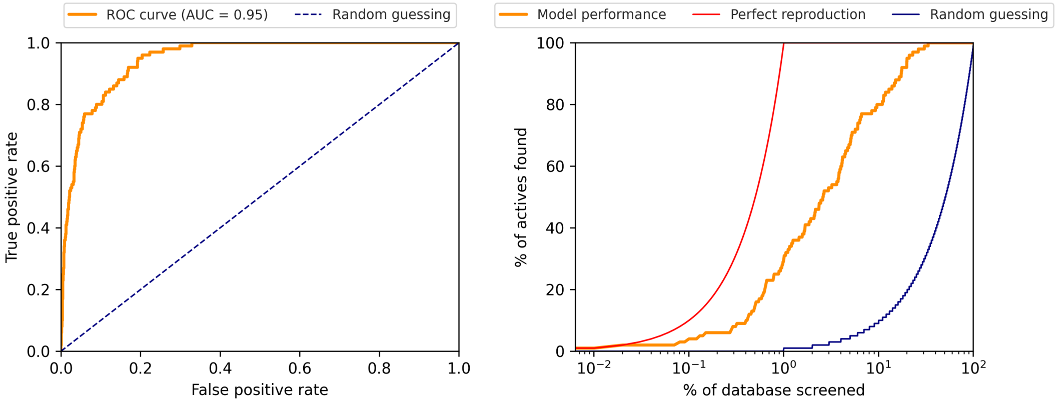


**Fig. S2**: Screening performance for query ZINC000950159323. Left: ROC curve, right: reproduction performance.

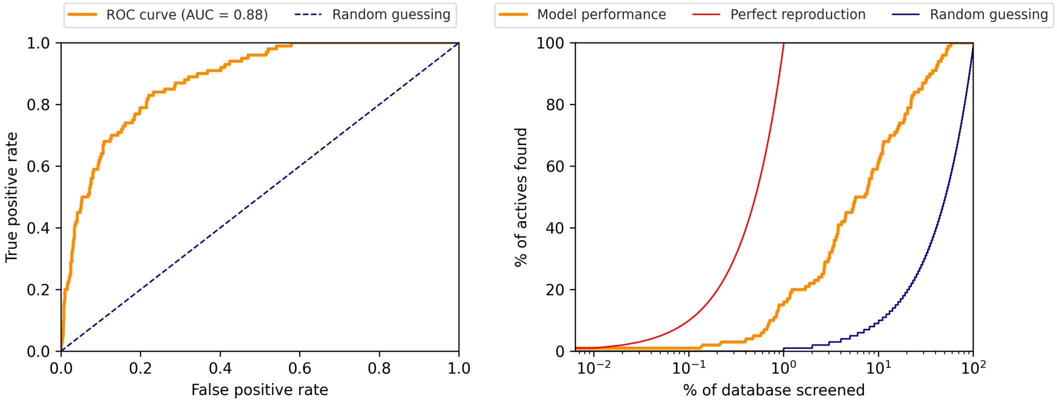


**Fig. S3**: Screening performance for query ZINC000954430177. Left: ROC curve, right: reproduction performance.

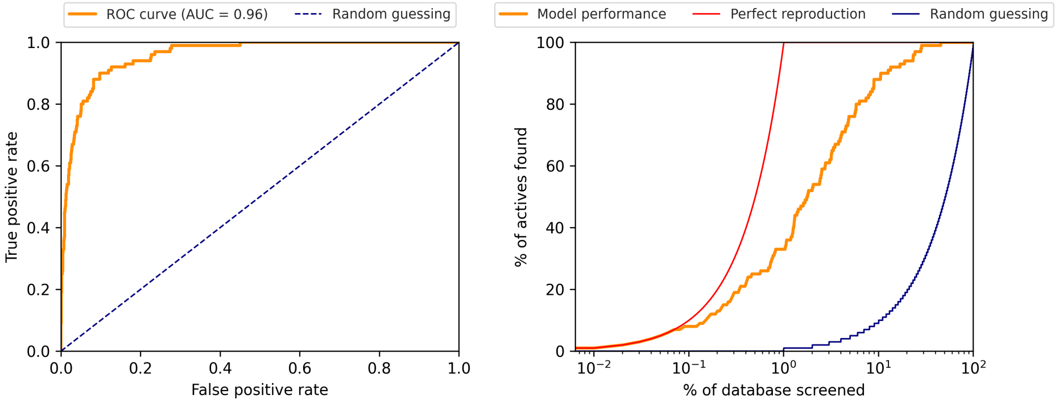


**Fig. S4**: Screening performance for query ZINC000970035445. Left: ROC curve, right: reproduction performance.

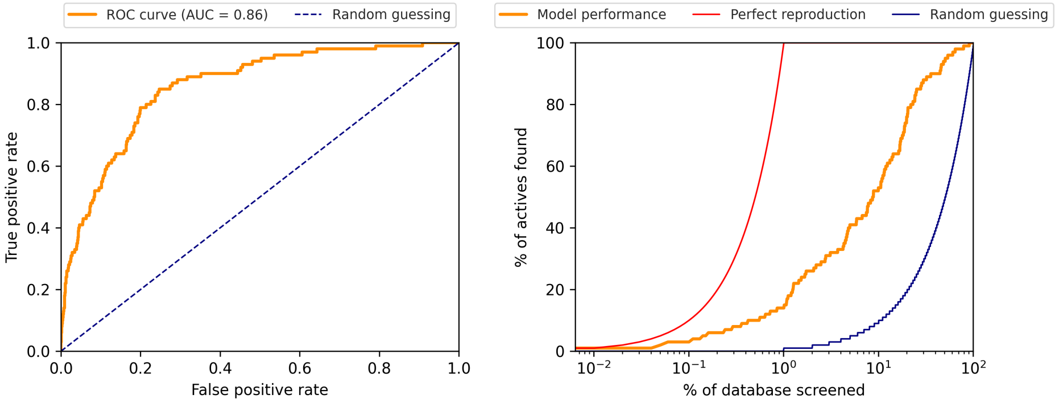


**Fig. S5**: Screening performance for query ZINC001183157671. Left: ROC curve, right: reproduction performance.

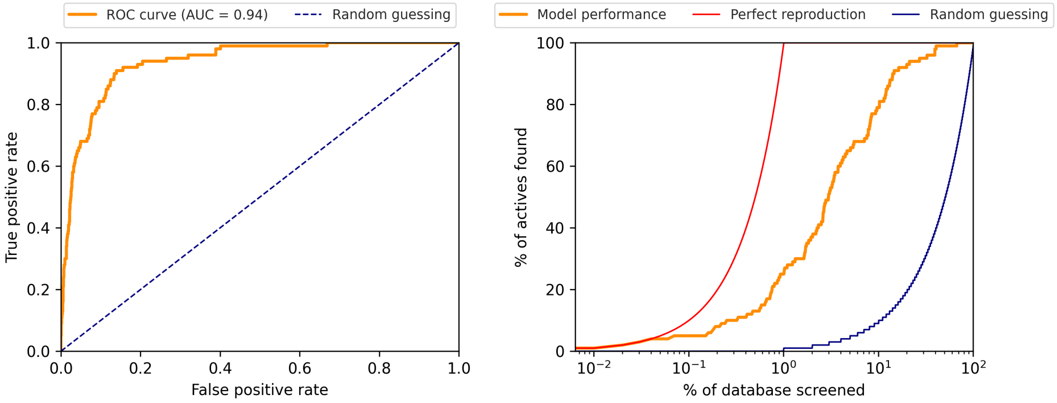


**Fig. S6**: Screening performance for query ZINC001281147597. Left: ROC curve, right: reproduction performance.

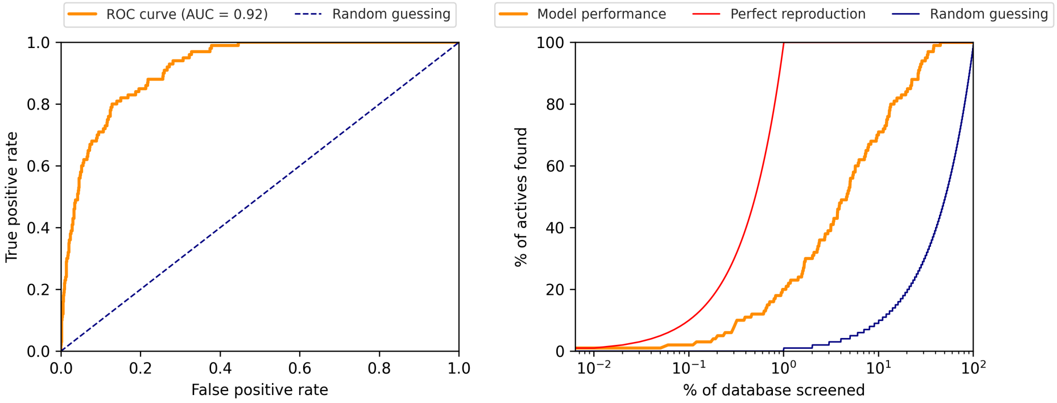


**Fig. S7**: Screening performance for query ZINC001368797027. Left: ROC curve, right: reproduction performance.

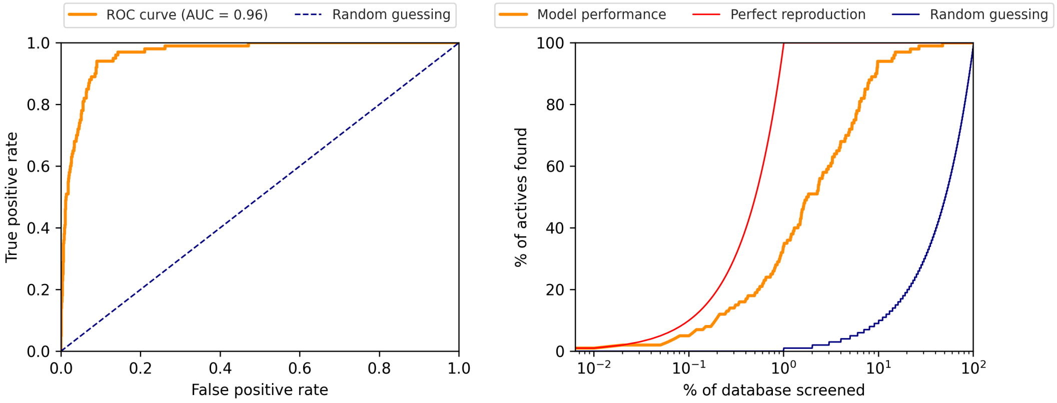


**Fig. S8**: Screening performance for query ZINC001711902206. Left: ROC curve, right: reproduction performance.

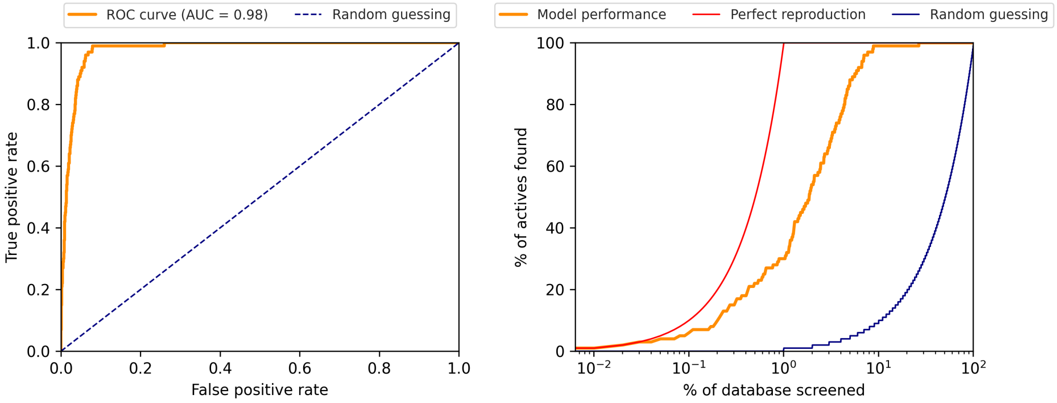


**Fig. S9**: Screening performance for query ZINC001740566933. Left: ROC curve, right: reproduction performance.

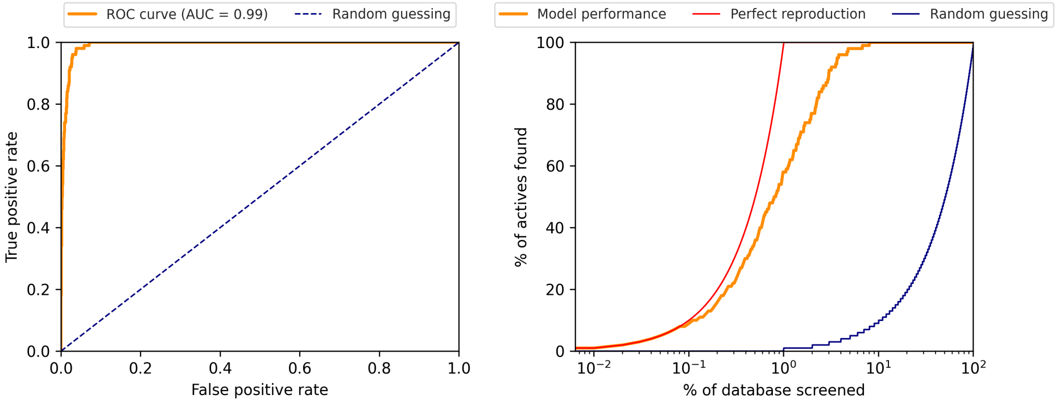


**Fig. S10**: Screening performance for query ZINC001763434742. Left: ROC curve, right: reproduction performance.

**Acknowledgments.** We gratefully acknowledge the support of NVIDIA Corporation with the donation of two RTX A5000 GPUs used for this research.
